# OM-85 reduces SARS-COV-2 viral RNA expression in nasopharyngeal cells from COVID-19 patients

**DOI:** 10.1101/2022.07.29.502045

**Authors:** Gisele Cassão, Krist Helen Antunes, João Ismael Budelon Gonçalvez, Leonardo Duarte Santos, Bruno Lopes Abbadi, Cristiano Valim Bizarro, Pablo Machado, Luiz Augusto Basso, Christian Pasquali, Renato T. Stein, Ana Paula Duarte de Souza

**Author notes:** These authors contributed equaly to this study (Gisele Cassão, Krist Helen Antunes).

## Abstract

OM-85 is a bacterial lysate from common respiratory tract pathogens, with an excellent safety profile, widely used to prevent recurrent respiratory tract infections. Several studies have been reporting the antiviral roles of OM-85. Here we demonstrated the effect of *ex-vivo* OM-85 exposure in nasopharyngeal cells collected from COVID-19 patients. OM-85 decreased the SARS-CoV-2 N1 gene expression and increased RIG-I (DDX58) gene expression in these cells. These data support the antiviral effect of OM-85 against SARS-CoV-2.

## 1. Introduction

The COVID-19 pandemic has affected a huge number of people around the world. Although the development of vaccines and the adoption of efficient and sanitary measures, several variants are still very active and affect individuals. There is an urgent need to develop new effective therapeutic approaches to minimize the burden of these severe respiratory viral infections and all associated sequelae. Treatments based on microbial-derived products, such as probiotics and postbiotics, can prevent and treat viral-induced respiratory diseases. Many of these studies have focused on the use of microbial therapies administered by the oral route, inducing the gut-lung axis into producing a potent host antiviral response. Recently, the delivery of these products through the airways is emerging as a safe and effective alternative strategy [1]. OM-85 is a postbiotic product containing a standardized lysate obtained by chemical lysis of 21 microorganisms strains associated with respiratory infections. OM-85 contains different microbial products, such as bacterial metabolites, proteins, peptides, traces of fatty amino acids, and saccharides. This bacterial lysate is widely used to prevent recurrent respiratory infection when administered orally [3, 4]. The protective effects of OM-85 are mainly due to its modulation of both cellular and humoral responses [3, 5, 6], presenting an anti-inflammatory effect and improving antiviral response mediated by interferons. OM-85 can be effective in preventing severe coronavirus infection in mice when administered through the oral route [7]. Additionally, the SARS-CoV-2 receptor ACE-2 is downregulated in the lungs of mice treated intranasally with OM-85 [8]. The concept that OM-85 directly administrated into the airways modulates immune response has been previously explored to prevent asthma and respiratory syncytial virus infection in mice [9, 10]. In this study, we aimed to evaluate the effect of *ex-vivo* treatment with OM-85 in nasopharyngeal cells isolated from COVID-19 patients.

## 2. Materials and Methods

We collected nasopharyngeal samples from adult hospitalized COVID-19 patients positive for SARS-CoV-2. Viral presence was confirmed by RT-PCR in the 48h before inclusion. Subject enrollment occurred from March to July 2021. The following variables were considered exclusion criteria: patients on mechanical ventilation; pregnant or breastfeeding women; patients with severe liver disease (alanine aminotransferase and/or aspartate aminotransferase > 5x normal); severe nephropathy (kidney transplantation or dialysis), HIV infection, cancer, hereditary angioedema, other immunodeficiencies, previous myocardial ischemic disease, previous thromboembolic disease. The cells were characterized staining with anti-CD45 (BD Bioscience) and analyzed by Flow cytometry. Also, the cells were centrifuged and stained with hematoxylin and eosin (Panotico rápido Laborclin).

To perform the *ex-vivo* treatment nasopharyngeal samples were centrifuged, washed with sterile PBS-EDTA (5mM) solution, and the cell pellet was suspended with Dulbecco’s Modified Eagle Medium (Nutrient Mixture F-12) (DMEM/F-12), 20mM HEPES (pH 7.4), 5% fetal bovine serum (FBS), and 0.06mM NaHCO3. Cells were seeded in 96-well-plates and treated with 10μg/ml OM-85 concentrate (OM-Pharma) or 100 ng/mL of human recombinant IFN-β protein (ThermoFisher Scientific). The concentration of 10μg/ml of OM-85 was based on previous data with RSV we recently generated [9]. After 24h, cells were collected for analysis by flow cytometry and the gene expression analysis by qRT-PCR. For viral gene expression we used 2019-nCoV N1 and RNaseP primers and probes (QuatroG, #100199). For cytokines expression we used TaqMan specific primers and probes (*ACTB* Hs03023943_g1; *IFNB1* Hs01077958_s1; *IL1B* Hs01555410_m1; *IFIH1* Hs00223420_m1; *DDX58* Hs01061436_m1; *ISG15* Hs01921425_s1; *OASL* Hs00984387_m1;

*IL6* Hs00174131_m1; *IL8* Hs00174103_m1; *TNFA* Hs00174128_m1. ThermoFisher Scientific). PCR conditions followed the TaqMan™ Universal PCR Master Mix protocols (Thermo Fisher Scientific, Waltham, MA, USA). Quantification of gene expression was conducted using StepOne™ (Applied Biosystems). The threshold cycle value (ΔCt) was obtained subtracting the Ct value from the endogenous gene by the Ct value of the target gene. The gene expression was calculated according to 2-^ΔCt^ formula. The supernatant was collected for cytokine analysis by ELISA (pbl Assay Science). We chose 24h of culture based on your previous study using the nasopharyngeal cells [11]. The study design was approved by the local ethical committee under the protocol CAAE n° 30754220.3.0000.5336. The data were summarized as mean and standard error of the mean (SEM). Univariate comparisons between groups were performed using the Mann-Whitney test, Wilcoxon test and Kruskal-Wallis for multiple comparisons (with Dunn’s multiple comparison test as post hoc). The data were analyzed using IBM SPSS Statistics Version 25.0 (IBM Corp, NY, USA) and GraphPad Prism Version 8.0 (GraphPad Software, CA, USA).

## 3. Results and discussion

Among the 28 patients enrolled to nasopharyngeal sample collection, there were 17 males (57%), at a median age of 48.96 years (CI 95% 44.49-53.43); mean body mass index (BMI) 31.94Kg/m^2^; and 64% were on oxygen supplementation. Table 1 describes the clinical and demographic characteristic of the patients that the nasopharyngeal samples were collected.

**Table 1.** Demographic and clinical characteristics of the patients (n = 28)

|  |  |
| --- | --- |
| *Gender, n (%)* |  |
| Male | 16 (57) |
| *Age, years old (mean ± SD)* | 48,96 ± 12,35 |
| *BMI, Kg/m<sup>2</sup> (mean ± SD)* | 31,94 ± 6,29 |
| *Axillary temperature, °C (mean ± SD)* | 36,01 ± 0,68 |
| *SPO<sub>2</sub>, % (mean ± SD)* | 95% ± 0,021 |
| *Oxygen use (mean, %)* | 18 (64) |
| *Corticoid use (mean, %)* | 25 (89) |
| *Antibiotic use (mean, %)* | 8 (29) |
| *Comorbidities, n* |  |
| Obesity | 11 |
| Systemic arterial hypertension (SAH) | 7 |
| No comorbidities | 6 |
| Asthma | 3 |
| Bacterial infection | 2 |
| Ex- smoker | 2 |
| Diabetes Mellitus (DM) | 2 |
| Hypothyroidism | 2 |
| Chronic obstructive pulmonary disease | 2 |
| Dyslipidemia | 2 |
| Rhinitis | 2 |
| Cardiovascular | 1 |
| Smoker | 1 |
| Vitiligo | 1 |
| Depression | 1 |
| Anxiety | 1 |
| Thromboembolism | 1 |

Definition of abbreviations: Kg/ m<sup>2</sup>, kilograms per square meter; SD, standard deviation; °C, degree Celsius.

Before treatment, nasopharyngeal cells were characterized and most of the population were composed by epithelial cells (Figures 1 a and b). We found that OM-85 *ex-vivo* treatment decreased the expression of SARS-CoV-2 spike protein on nasopharyngeal cells from COVID-19 patients (Figure 1c).

**Figure 1.**
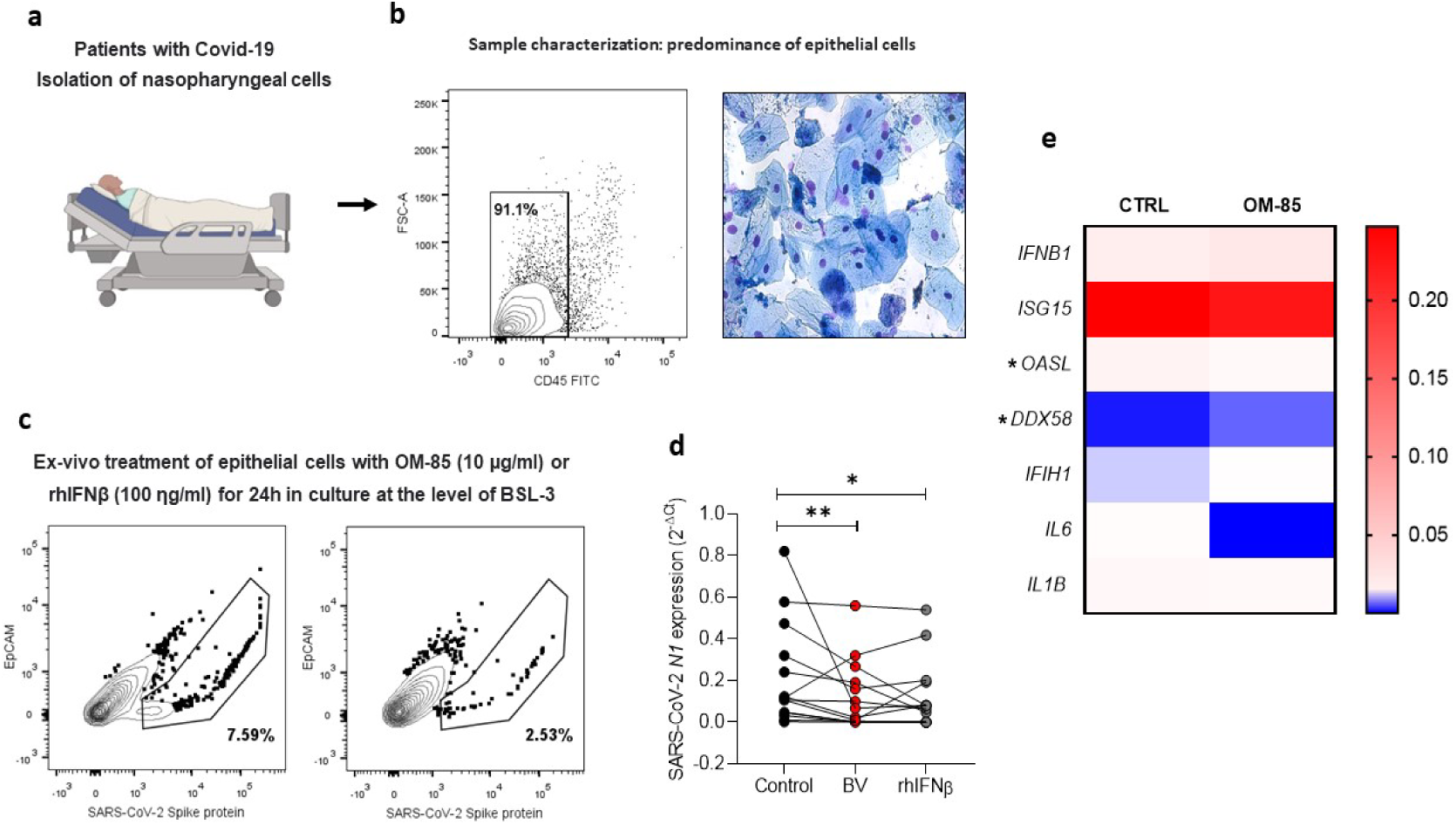
OM-85 *ex-vivo* exposure on fresh isolated nasopharyngeal cells from COVID-19 patients decrease SARS-CoV-2 viral gene expression. (A) Experimental design: Nasopharyngeal lavage samples were processed to obtain respiratory epithelial cells for ex vivo analysis to investigate the effect of OM-85 treatment. b Characterization of nasopharyngeal cells. Most of cells (91.1%) are CD45 negative and presented characteristic of epithelial cells after cytocentrifuged and stained c Cells were cultured for 24 hours in the presence of OM-85 (10 μg /ml) and recombinant IFNβ (100 ng/ml) and stained with anti-SARS-CoV-2 spike protein and anti-EpCam. d Expression of the SARS-CoV-2 N1 gene by real-time PCR. The difference between the treatment and its untreated control was assessed by the Wilcoxon matched-pairs signed-rank test (*p<0.05, **p<0.01). e Heatmap showing the data from gene expression (*p < 0.05).

These data are in line with the previously seen effect of OM-85 pre-treatment on SARS-CoV-2 infection (8). Interestingly, expression of viral N1 gene was significantly reduced in cells treated with OM-85 compared to untreated control (Figure 1d). The effect of OM-85 on SARS-CoV-2-infected human nasopharyngeal cells e*x-vivo* was similar to hrINF-β̲treated cells (Figure 1d). To further examine these findings, we next analyzed the expression of antiviral response-associated genes, and their corresponding differential expression are shown in Figures 1e. OM-85 did neither modulated the expression of IFN-β (n=12; p=0.3804), nor the responsive genes ISG15 (n=26; p=0.7454) and MDA-5 (n=12; p=0.5186). Considering its importance in the context of COVID-19, the RIG-I viral recognition pathway was also investigated and here, OM-85 treatment significantly augmented RIG-I expression (n=8; p=0.0156). In contrast to RIG-I, OASL (n=19; p=0.0289) had its expression significantly reduced after OM-85 treatment. More in-depth analysis on soluble mediators and endotypes revealed no significant difference between the OM-85 treated and control groups. These was particularly true for genes associated with the inflammatory response, i.e., IL-6 (n=5; p=0.0625) and IL1-β (n=18; p=0.737). Gene expression of ACE-2, IL-8, MAVS, IRF-3, mTOR, IFN-λ, TRL2 and TLSP, were not detected in the samples, possibly due to the low expression of these genes by the time of harvesting. We did not find differences on cytokine levels in the supernatant between untreated and treated cells. (Table 2).

**Table 2:** Cytokine analysis in the supernatant of the treated nasopharyngeal cells

| Cytokine | Median<br>Control | Median<br>OM-85 | p |
| --- | --- | --- | --- |
| IFN-β (pg/ ml) n= 9 | 575,9 | 809,3 | 0,6665 |
| IL-6 (pg/ ml) n= 8 | 4,451 | 2,177 | 0,5041 |
| IL-8 (pg/ ml) n=21 | 2062 | 2073 | 0,8813 |
| IL-10 (pg/ ml) n= 10 | 13,98 | 18,27 | 0,5639 |
| TNF-α (pg/ ml) n=10 | 15,50 | 24,67 | 0,4692 |

(Mann-Whitney test)

Therefore, while OM-85 did not significantly modulate the production of IFN-β, among other cytokines, interestingly there was an increase in the expression of RIG-I and reduced the expression of OASL. We postulate this might be associated with different cell signaling pathways involved with the time until OM-85 treatment, clinical characteristics, and individual immune responses. These aspects need to be further investigated in future studies.

One strength of our study is that we tested the antiviral effect of OM-85 directly on airway cells freshly collected from COVID-19 patients. The epithelium in the upper respiratory tract is a frontline barrier in the battle against SARS-CoV-2, several studies have highlighted the importance of the analysis of samples from the airways to better characterize the direct mucosal impact of possible interventions on SARS-CoV-2 infection [12-14]. In spite these findings, our study suffers from two limitations, one is associated with the number of cells obtained from the patients, limiting the analysis to a one-time point, and the second is the lack of the evaluation of OM-85 effect on virus replication on these cells. Although, studies have demonstrated that the virus can replicate on the nasal epithelial cells [15] and after 24h of infection in cell lines [16].

Considering the former, our results still identified that RIG-I activation might be one of the pathways related to the OM-85 effect on nasopharyngeal cells during SARS-CoV-2 infection. Antiviral responses associated with RIG-I activation are essential to control SARS-CoV-2 infection [17]. Nevertheless, SARS-CoV-2 proteins have been described to inhibit antiviral responses by interacting with RIG-I [18, 19]. Efficient antiviral activity in the airway should point to a balanced immune response leading to viral clearance and faster patient recovery. Severe viral respiratory infection is related to an aberrant immune response leading to uncontrolled inflammation. The role of type 1 interferon in COVID-19 severity is controversial [20]. Although a large study conducted by WHO did not show clinical benefit of subcutaneous interferon beta 1-a treatment on hospitalized COVID-19 patients [21], a phase 2 trial showed a promising result of inhaled nebulized interferon beta-1a [22]. Another intervention tested by the inhalation route to COVID-19 patients is budesonide [23].

In summary, we identified that OM-85 can have an antiviral effect on SARS-CoV-2 infected nasopharyngeal cells collected from COVID-19 patients. These findings highlight a potential therapeutic effect of OM-85 against viral infection on the respiratory tract, a tissue that could be directly accessed in subjects using the intranasal route. Proper human intervention studies are needed to confirm these findings so that OM-85 becomes an alternative viable therapy by intranasal route for patients with respiratory infections.

## Author Contributions

Conceptualization, A.P.D.S and R.S.; formal analysis, G.C, K.H.A, J.I.B.G., M.D; B.L.A., investigation, G.C, B.L.A, L.D.S., K.H.A, M.D. and J.I.B.G.; resources, P.M, C.V.B, L.A.B; writing—original draft preparation, G.C and K.H.A.; writing—review and editing, A.P.D.S, C.P, R.S.; supervision, A.P.D and R.S.; project administration, A.P.D.S .; funding acquisition, A.P.D.S, C.V.B, L.A.B, P.M and R.S. All authors have read and agreed to the published version of the manuscript.

### Funding

This study was funded by FAPERGS COVID-19 20/2551-0000258-6 and Coordenação de Aperfeiçoamento de Pessoal de Nível Superior - Brasil (CAPES) -Finance Code 001.

### Institutional Review Board Statement

The study was conducted in accordance with the Declaration of Helsinki and approved by the Ethics Committee of PUCRS (CAAE n° 30754220.3.0000.5336.l).

### Informed Consent Statement

Written informed consent has been obtained from the patient(s) to publish this paper.

### Data Availability Statement

The data presented in this study are not publicly available but are available upon request from the corresponding author.

## Acknowledgments

We thank Eduardo Pedraza for technical support.

